## Supplementary figures and images for "FUS regulates RAN translation through modulating the G-quadruplex structure of GGGGCC repeat RNA in *C9orf72*-linked ALS/FTD"

### Actin raw.jpg

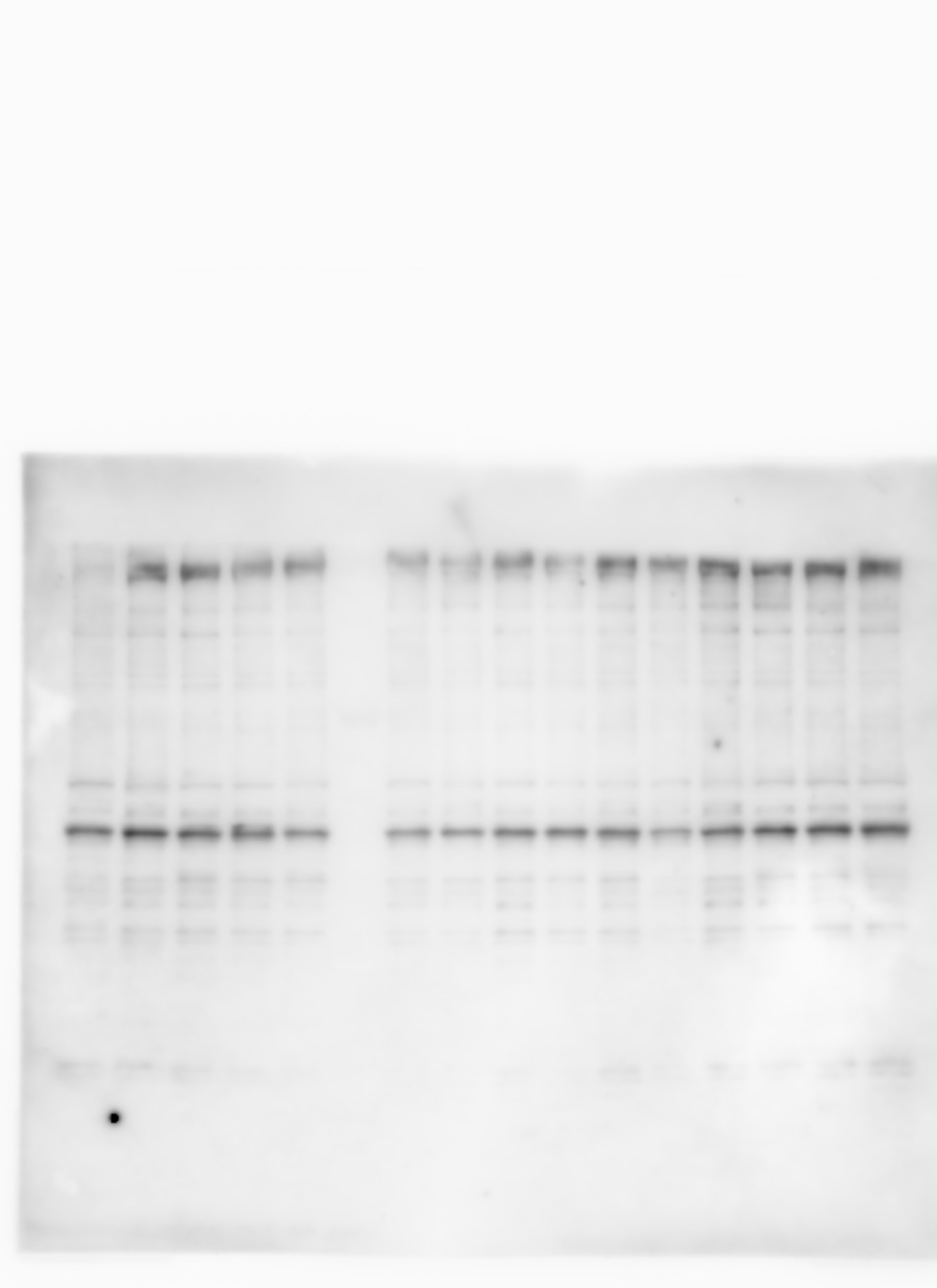

### Actin with labels.jpg

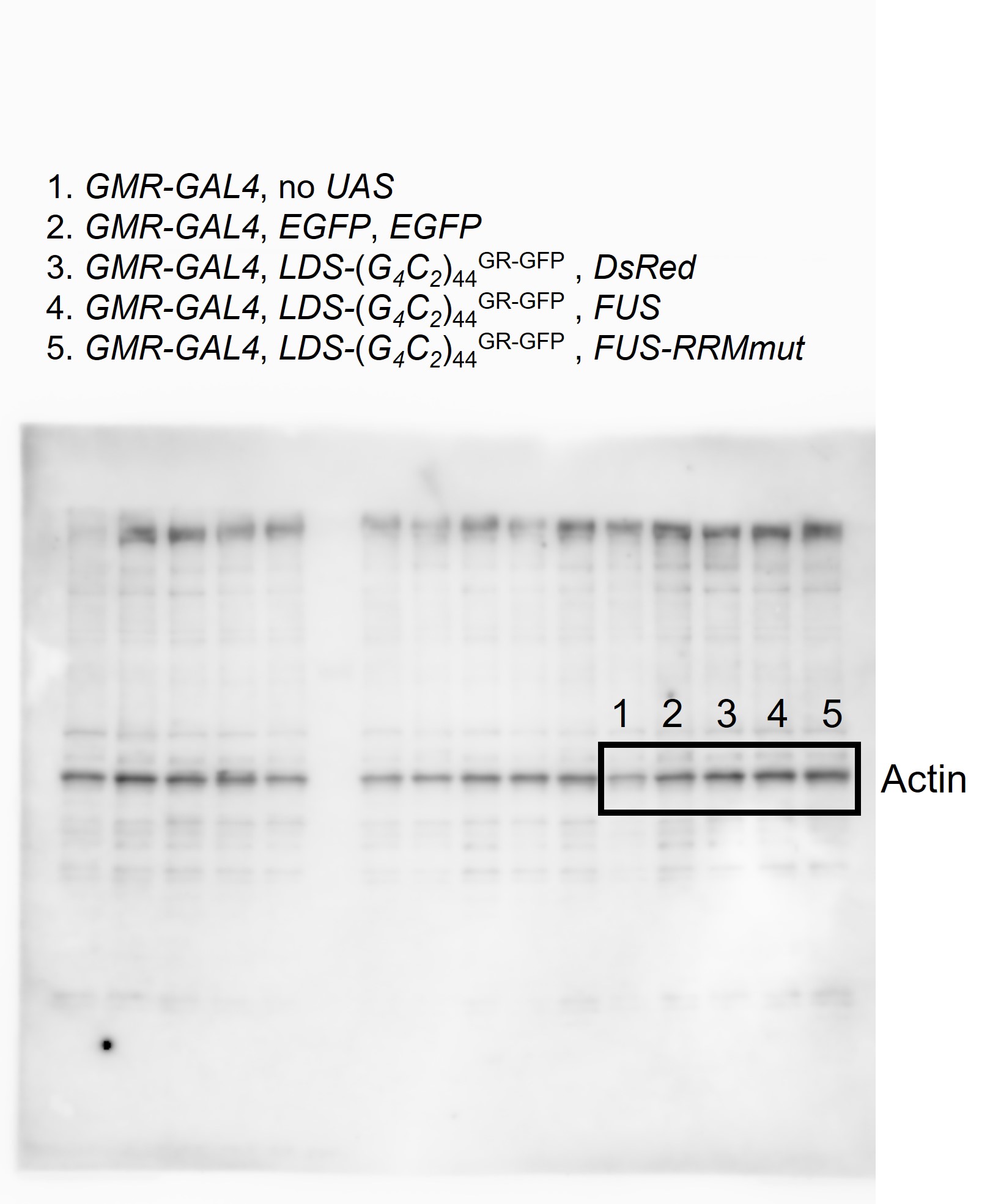

### EGFP raw.jpg

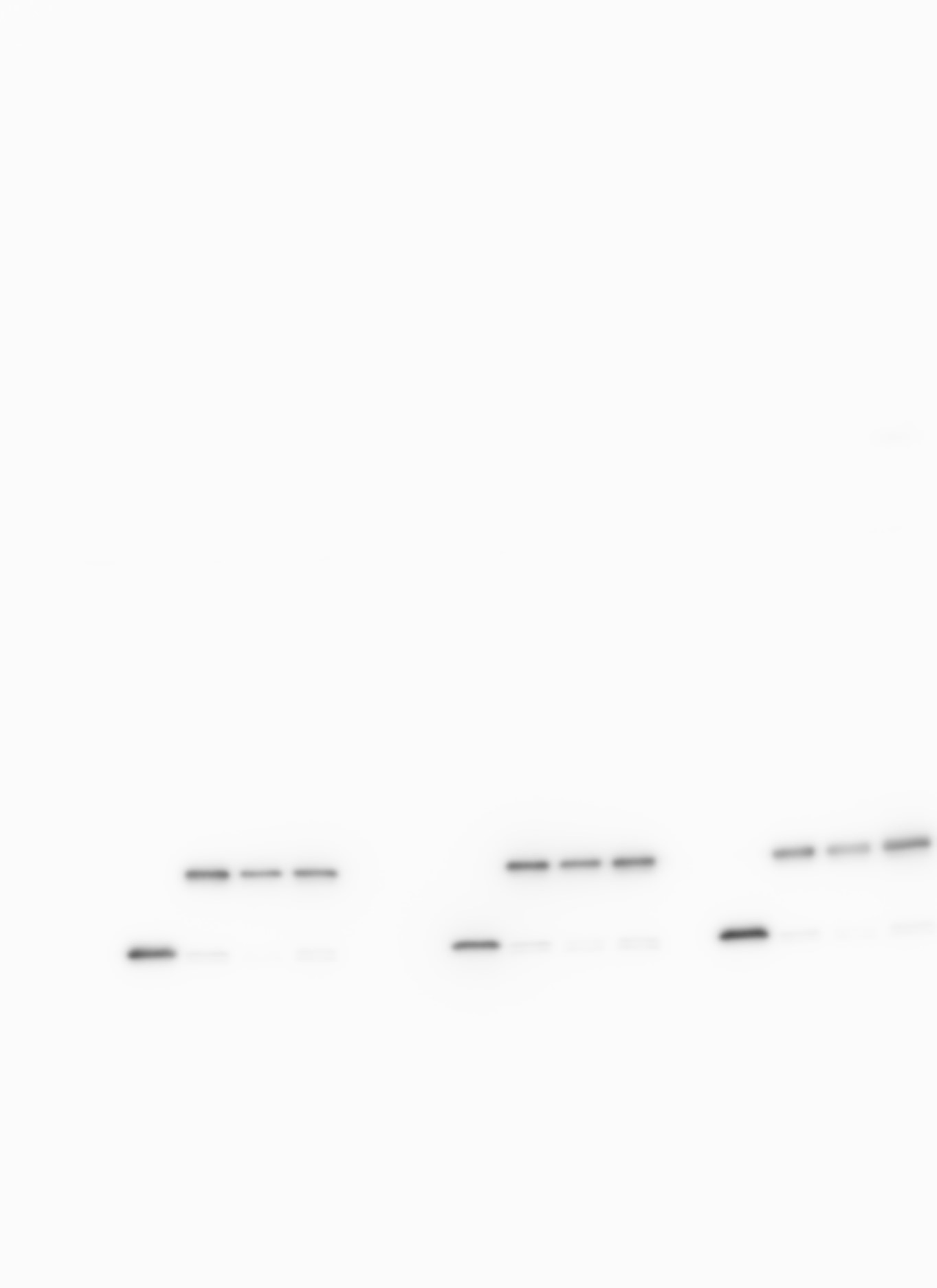

### EGFP with labels.jpg

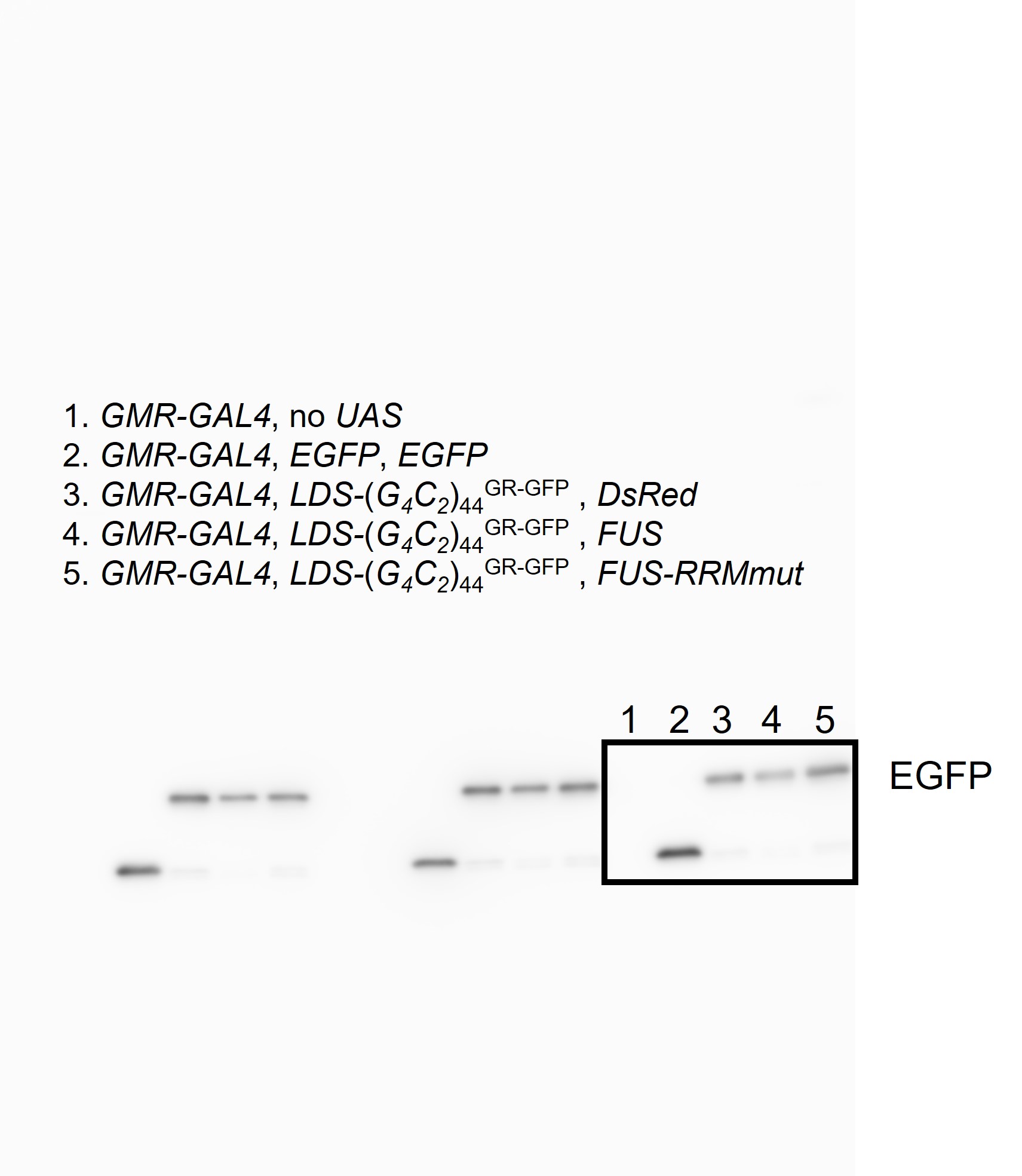

### FUS and action raw.jpg

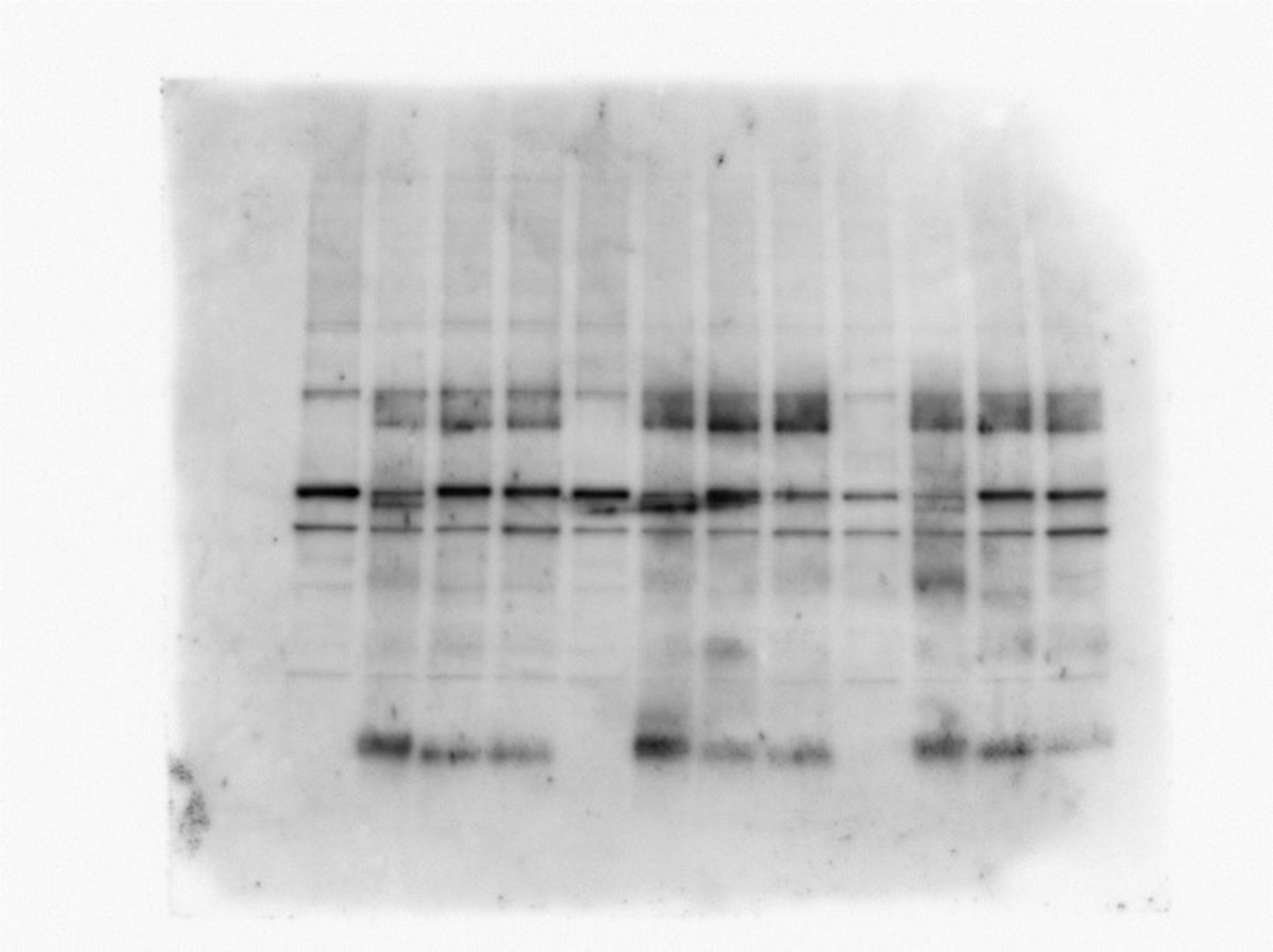

### FUS and action with labels.jpg

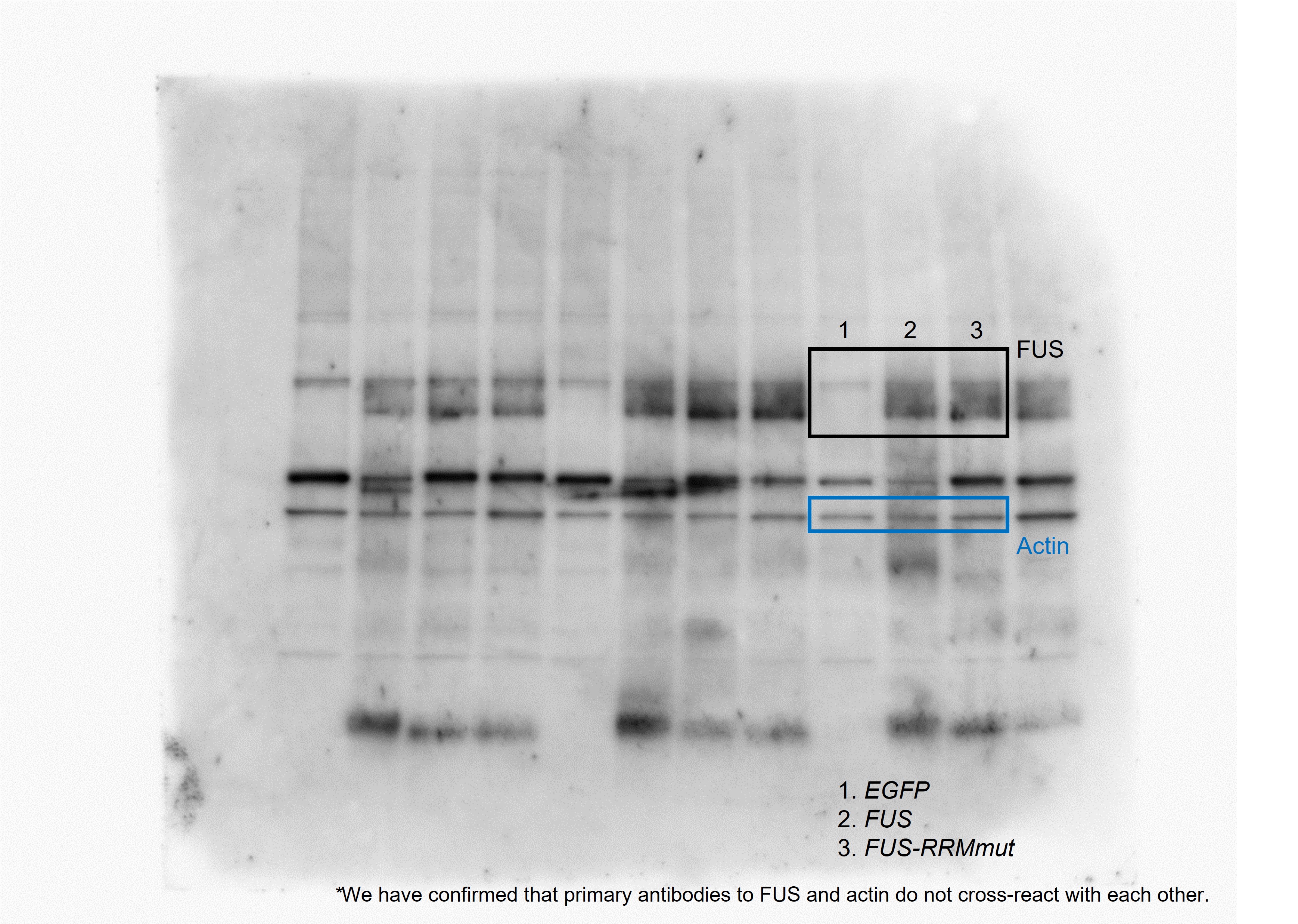

### GA-Myc raw.jpg

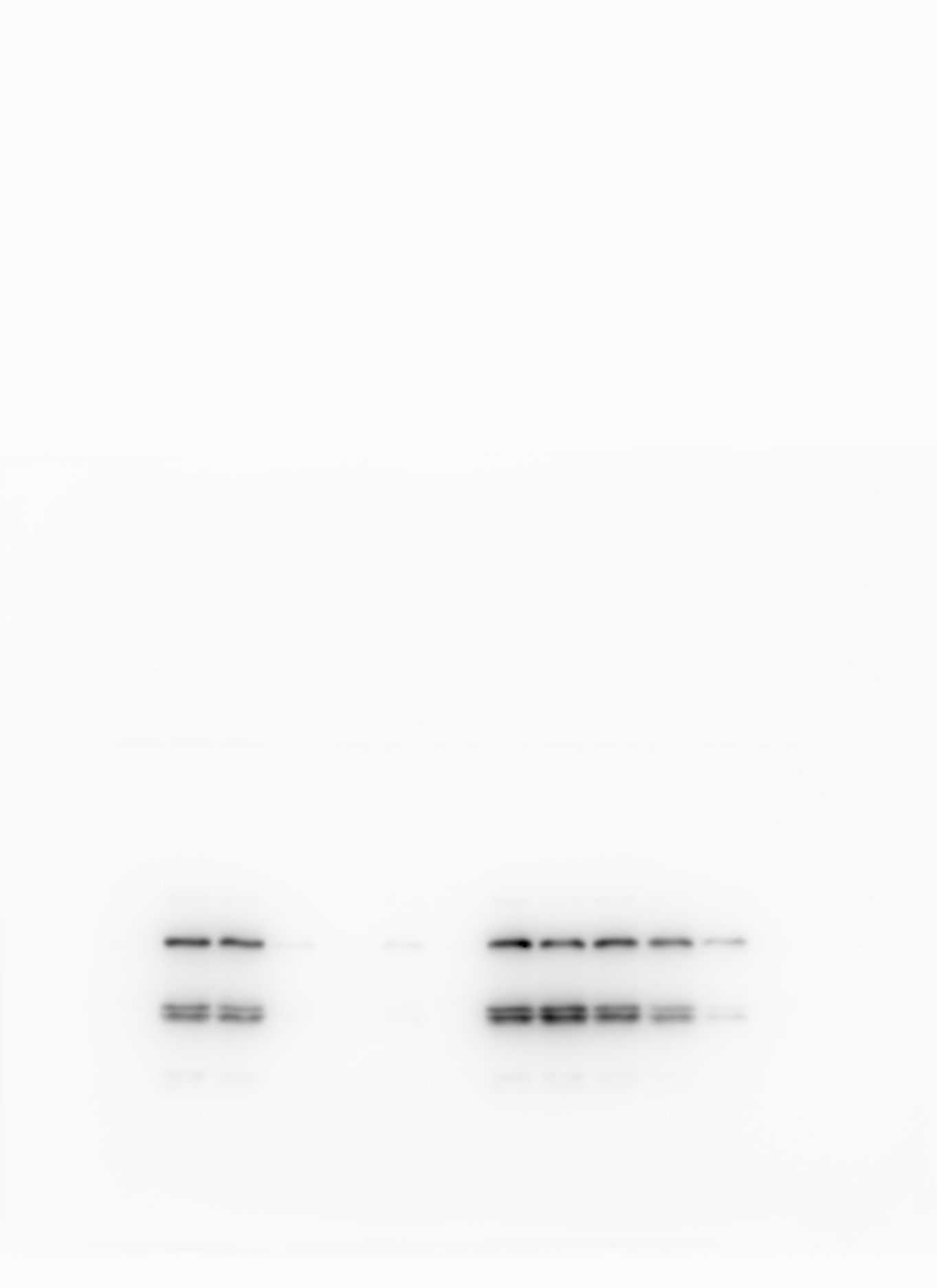

### GA-Myc with labels.jpg

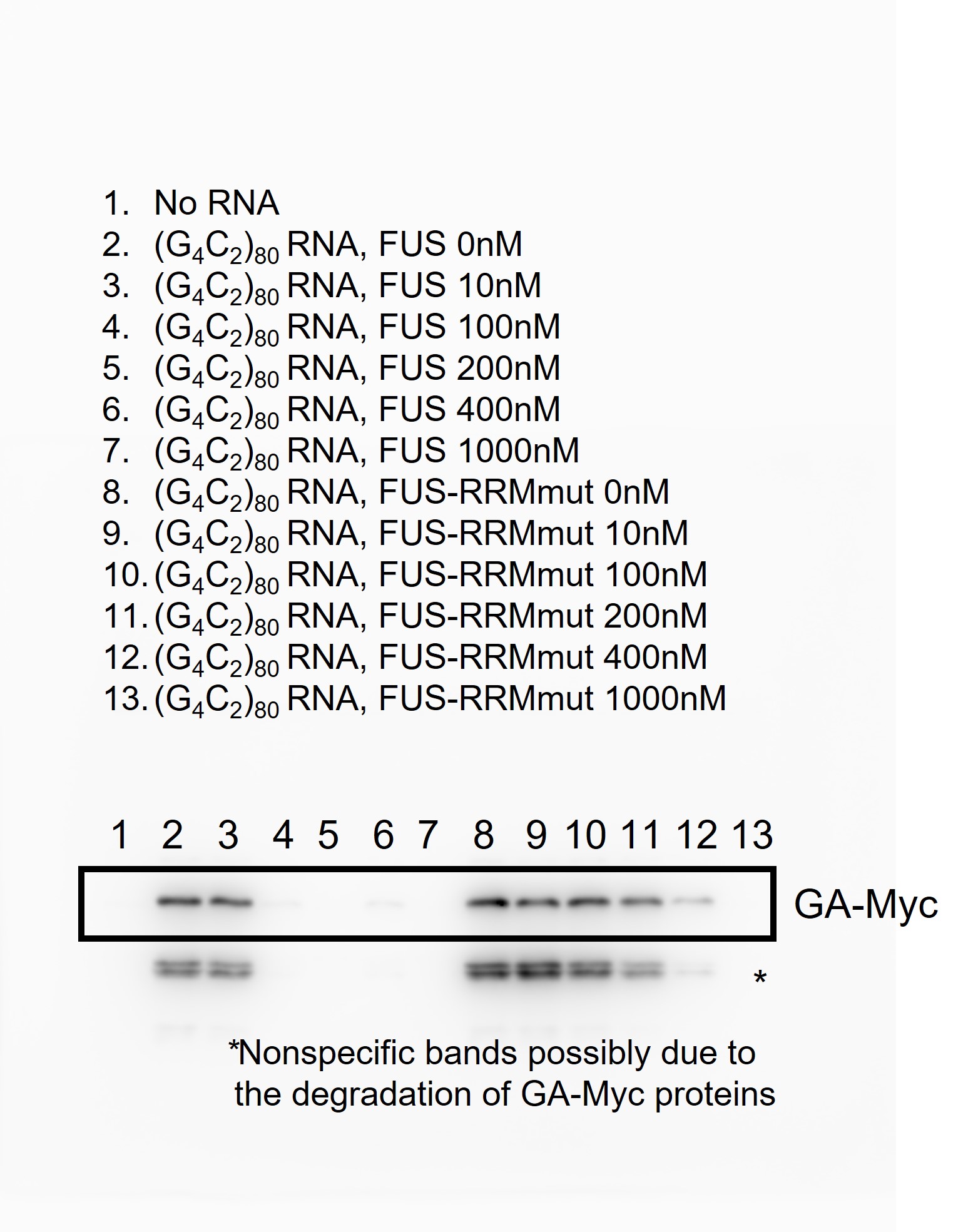

### GR raw.jpg

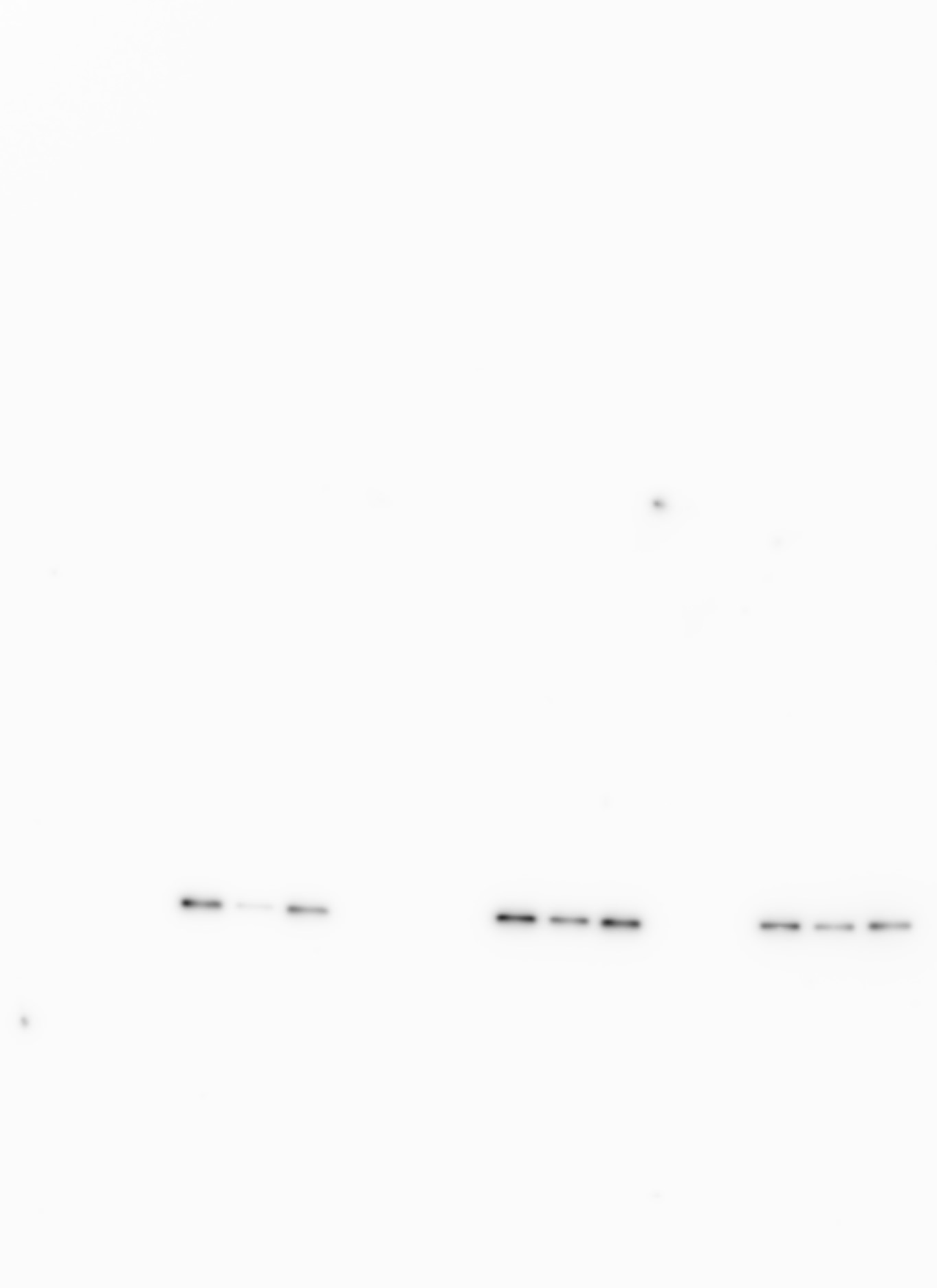

### GR with labels.jpg

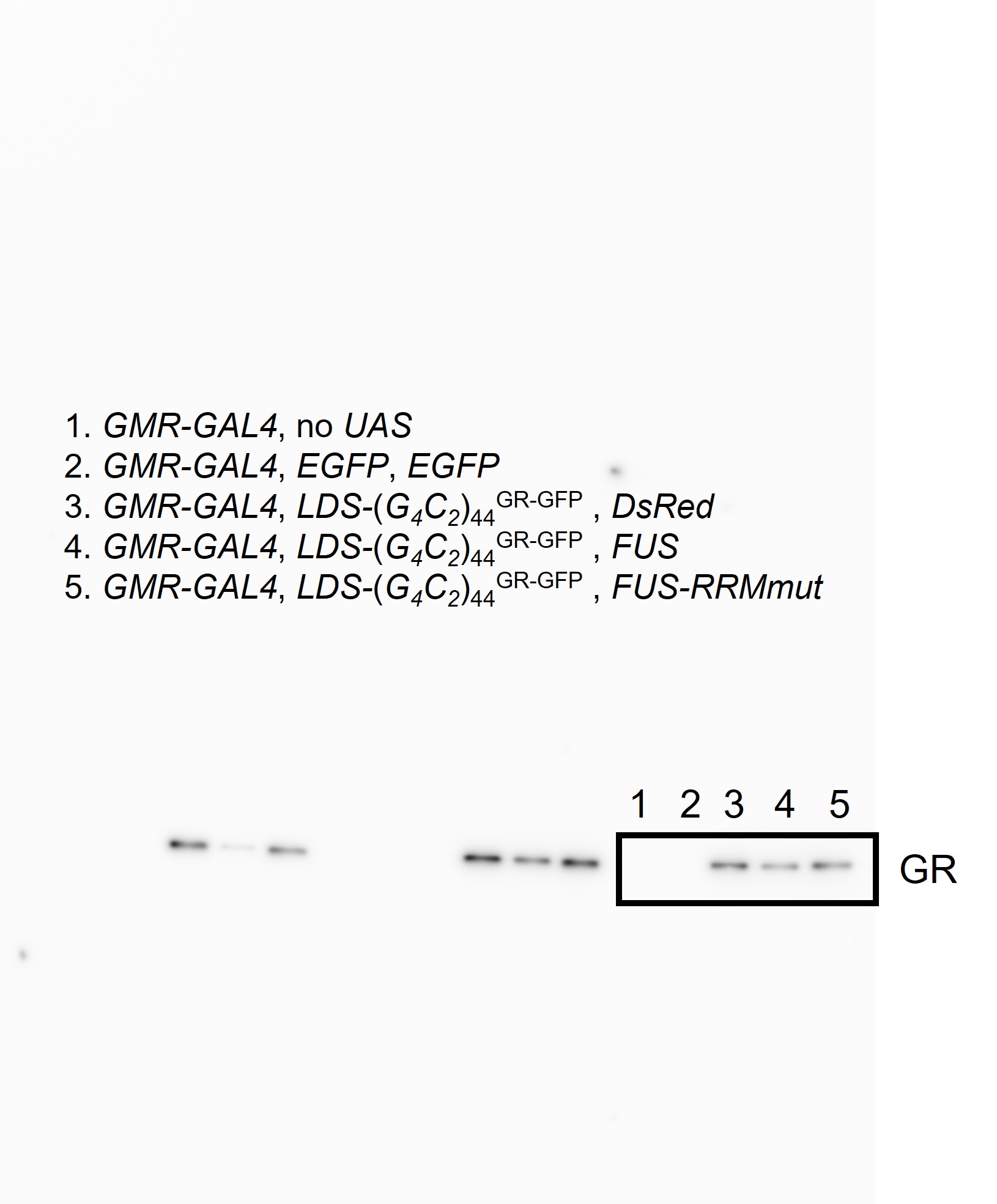
